# Accuracy of genomic selection for growth and wood quality traits in two control-pollinated progeny trials using exome capture as genotyping platform in Norway spruce

**DOI:** 10.1101/293696

**Authors:** Zhi-Qiang Chen, John Baison, Jin Pan, Bo Karlsson, Bengt Andersson Gull, Johan Westin, María Rosario García Gil, Harry X. Wu

## Abstract

**Background:** Genomic selection (GS) can increase genetic gain by reducing the length of breeding cycle in forest trees. Here we genotyped 1370 control-pollinated progeny trees from 128 full-sib families in Norway spruce (*Picea abies* (L.) Karst.), using exome capture as a genotyping platform. We used 116,765 high quality SNPs to develop genomic prediction models for tree height and wood quality traits. We assessed the impact of different genomic prediction methods, genotype-by-environment interaction (G×E), genetic composition, size of the training and validation set, relatedness, and the number of SNPs on the accuracy and predictive ability (PA) of GS.

**Results:** Using G matrix slightly altered heritability estimates relative to pedigree-based method. GS accuracies were about 11–14% lower than those based on pedigree-based selection. The efficiency of GS per year varied from 1.71 to 1.78, compared to that of the pedigree-based model if breeding cycle length was halved using GS. Height GS accuracy decreased more than 30% using one site as training for GS prediction to the second site, indicating that G×E for tree height should be accommodated in model fitting. Using half-sib family structure instead of full-sib led a significant reduction in GS accuracy and PA. The full-sib family structure only needed 750 makers to reach similar accuracy and PA as 100,000 markers required for half-sib family, indicating that maintaining the high relatedness in the model improves accuracy and PA. Using 4000–8000 markers in full-sib family structure was sufficient to obtain GS model accuracy and PA for tree height and wood quality traits, almost equivalent to that obtained with all makers.

**Conclusions:** The study indicates GS would be efficient in reducing generation time of a breeding cycle in conifer tree breeding program that requires a long-term progeny testing. Sufficient number of trees within-family (16 for growth and 12 for wood quality traits) and number of SNPs (8000) are required for GS with full-sib family relationship. GS methods had little impact on GS efficiency for growth and wood quality traits. GS model should incorporate G × E effect when a strong G×E is detected.

## Background

Norway spruce (*Picea abies* (L.) Karst.) is one of the most important conifer species for commercial wood production and ecological integrity in Europe [1]. A conventional breeding program for Norway spruce based on pedigree-based phenotypic selection usually takes between 20–30 years in Scandinavian countries [2]. To shorten the breeding cycle, genomic selection (GS) has recently been proposed as an alternative in many tree species such as eucalypts (*Eucalyptus*) [3-5], maritime pine (*Pinus pinaster* Aiton) [6, 7], loblolly pine (*Pinus taeda* L.) [8, 9], white spruce and its hybrids (*Picea glauca* [Moench] Voss) [10-13], and black spruce (*Picea Mariana* [Mill] B.S.P.) [14].

Hayes et al. [15] considered four major factors affecting the accuracy of GS: 1) heritability of the target trait; 2) the extent of linkage disequilibrium (LD) between the marker and the quantitative trait locus (QTL); 3) the size of training population and the degree of relationship between training set (TS) and validation set (VS); and 4) the genetic architecture of the target trait. The genetic architecture and heritability of the target/breeding traits are intrinsic of the nature of the traits and environment in the test trials. Thus it is difficult to change in a practical breeding process. However, some factors such as LD between the marker and the QTL and the size of training population are relatively easy to be managed by increasing the number of markers and relationship between training and prediction (or validation) populations, reducing effective population size (*N*_*e*_), and increasing training population size [16].

Model selection in GS is quite important for the prediction of genomic estimated breeding value (GEBV) [15]. In GS, model adequacy is more related to genetic architecture of the target trait. Based on genome-wide association studies (GWAS), most growth and wood quality traits for conifer species have polygenic inheritance with a gamma or exponential distribution of allelic effects [10, 17]. To account for these skewed distributions of a few genes with large effects and most of genes with small effects, Bayes A, B, and Cπ and Bayesian Least Absolute Shrinkage and Selection Operator (BLASSO) were developed to fit the models more accurately, in contrast to Genomic best linear unbiased prediction (GBLUP) model that assumes a normal distribution of allelic effect. In most studies for growth and wood quality traits, the results were similar regardless of models used [4, 9]. Resende et al. [9] reported that fusiform rust in loblolly pine may be controlled by a few large genes and Bayesian based models had higher predictive ability (PA) defined as the correlation between adjusted phenotype value and GEBV. Thus, it is worthwhile to test different models on different traits which may have different genetic architectures for specific tree species.

So far, several genotyping technologies have been employed in GS, such as diversity array technology (DArT) array [5, 18], SNP chip/array [4-6, 9, 13, 14], genotyping by sequencing (GBS) [12, 13], and exome capture [19]. Those technologies were developed to genotype a subset of a whole genome, especially for conifer species with a large genome size. From the published papers, the number of used makers in tree species varies from 2500 to 69,511 single nucleotide polymorphisms (SNPs), most with a few thousands of SNPs. With the large genome size in most commercial conifer species (~ 20Gb) [20], for example, 20 Gb in Norway spruce [21], such small number of markers may not be able to capture most of QTL effects with short-range marker-QTLs LD in undomesticated populations or large breeding populations. Thus, such studies mostly capture those QTL effects with long-range maker-QTLs LD and relationships in highly related populations, such as full-sib families in tree breeding programs with a small *N*_*e*_.

Evaluations of accuracy in GS have been performed with phenotypic and dense maker data from a single site and multiple sites in several tree species [8, 12, 18]. Substantial G×E effects for growth traits have been found in conifer species [22-24]. However, wood quality traits usually have a low or non-significant G×E [25-27]. Thus, a GS model for growth traits based on a single site used to predict genomic breeding values in another site, may produce a low accuracy.

The aims of this study were to 1) evaluate the accuracy of GS on tree height and wood quality traits; 2) assess the GS accuracy for single site, cross-site, and joint-sites (e.g. G×E effect on GS selection); 3) examine effect of different statistical models and ratios between TS and VS for GS; 4) explore the roles of relatedness (full-sib, half-sib and unrelated) on accuracy of GS; 5) test the accuracy using subsets of random markers and markers with the largest positive effects; 6) estimate number of trees within-family and number of families required for effective GS for tree height and wood quality traits.

## Materials and methods

### Sampling of plant material

In this study, 1,370 individuals were selected from two 28-year-old control-pollinated progeny trials with 128 families from a partial diallel mating design that consisted of 55 parents originating from Northern Sweden. Buds and the first year fresh needles from 46 parents were sampled in a grafted archive at Skogforsk, Sävar (63.89°N, 20.54°E) and in a grafted seed orchard at Hjssjö (63.93°N, 20.15°E). Progenies were raised in the nursery at Sävar, and the trials were established in 1988 by Skogforsk in Vindeln (64.30°N, 19.67°E, altitude: 325 m) and in Hädanberg (63.58°N, 18.19°E, altitude: 240 m).

A completely randomized design without designed pre-block was used in the Vindeln trial (site 1), which was divided into 44 post-blocks. Each rectangular block has 60 trees (6×10) with expected 60 families at spacing of 1.5 m × 2 m. The same design was also used in the Hädanberg trial (site 2) with 44 post-blocks, but for the purpose of demonstration, there was an extra design with 47 extra plots, each plot with 16 trees (4×4). Based on the spatial analysis, in the final model, 47 plots were combined into two big post-blocks.

### Phenotyping

Tree height was measured in 2003 at the age of 17 years. Solid-wood quality traits including Pilodyn penetration (Pilodyn) and acoustic velocity (velocity) were measured in October, 2016. Surrogate wood density trait was measured using Pilodyn 6J Forest (PROCEQ, Zurich, Switzerland) with a 2.0 mm diameter pin, without removing the bark. Velocity is highly related to microfibril angle (MFA) in Norway spruce [28] and was determined using Hitman ST300 (Fiber-gen, Christchurch, New Zealand). By combining the Pilodyn penetration and acoustic velocity, indirect modulus of elasticity (MOE) was estimated using the equation developed in Chen et al. [28].

#### Genotyping

Total genomic DNA was extracted from 1, 370 control-pollinated progeny and their 46 unrelated parents using the Qiagen Plant DNA extraction protocol with DNA quantification performed using the Qubit^®^ ds DNA Broad Range Assay Kit, Oregon, USA. Probe design and evaluation is described in Vidalis et al. (2018, unpublished). Sequence capture was performed using the 40 018 probes previously designed and evaluated for the materials (Vidalis et al 2018, unpublished) and samples were sequenced to an average depth of 15x at an Illumina HiSeq 2500 platform. Raw reads were mapped against the *P. abies* reference genome v1.0 using BWA-mem [29, 30]. SAMTools [31] and Picard [32] were used for sorting and removal of PCR duplicates and the resulting BAM files were subsequently reduced to containing the probe only bearing scaffolds (24919) before variant calling. Variant calling was performed using Genome Analysis Toolkit (GATK) HaplotypeCaller [32] in Genome Variant Call Format (gVCF) output format. Samples were then merged into batches of ~200 before all samples were jointly called.

As per the recommendations from GATK’s best practices, Variant Quality Score Recalibration (VQSR) method was performed in order to avoid the use of hard filtering for exome/sequence capture data. The VQSR method utilizes machine-learning algorithms to learn from a clean dataset to distinguish what a good versus bad annotation profile of variants for a particular species should be like. For the VQSR analysis two datasets were created, a training subset and the final input file. The training dataset was derived from a Norway spruce genetic mapping population with loci showing expected segregation patterns (Bernhardsson et al. 2018, unpublished). The training dataset was designated as true SNPs and assigned a prior value of 12.0. The final input file was derived from the raw sequence data using GATK best practices with the following parameters: extended probe coordinates by +100 excluding INDELS, excluding LowQual sites, and keeping only bi-allelic sites. The following annotation parameters QualByDepth, MappingQuality and BaseQRankSum, with tranches 100, 99.9, 99.0 and 90.0 were then applied to the two files for the determination of the good versus bad variant annotation profiles.

The recalibrated Variant Call Format was filtered based on the following steps: (1) removing indels; (2) keeping only biallelic loci; (3) treating genotype with a genotype quality (GQ) < 6 as missing; (4) filtering read depth (DP) < 2; (5) removing individual call rate < 50%; (6) removing variant call rate (“missingness”) < 90%; (7) minor allele frequency (MAF) < 0.01. After steps 1−4, we calculated discordance between 148 pairs technique replicates, the average of discordance was less 1%. Thus, conditions of GQ <6 and DP < 2 as missing are sufficient to do downstream analysis. After all filtering, 116,765 SNPs were kept for downstream analysis.

LD K-nearest neighbor genotype imputation approach [33] was used to impute missing genotypes in TASSEL 5 [34]. After several rounds of imputation, a few of missing genotypes were imputed using random imputation of the codeGeno function in synbreed package in R [35].

### Estimating breeding values

Breeding values for the genotypes in the trials were predicted using the following model:

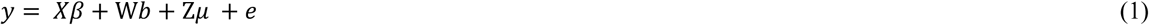

Where ***y*** is a vector of phenotypic observations of a single trait; *β* is a vector of fixed effects, including a grand mean and site effect, *b* is a vector of post-block within site effect, *μ* is a vector of site by additive effects of individuals. X, W, and Z are incidence matrices for *β*, *b*, and *μ*, respectively. For join-site cross-validation, the average of breeding values was assumed as estimated (true) breeding values (EBV). Otherwise, EBV in single site was assumed as true or reference breeding values. The random additive effects (*μ*) in equation (1) were assumed to follow 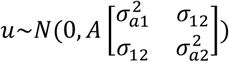, where *A* is the additive genetic relationship matrix, 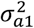 and 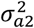 are the additive genetic variances for site 1 and site 2, respectively, *σ*_12_ is additive genetic covariance between site 1 and site 2. The residual *e* was assumed to follow 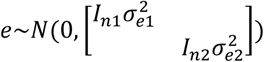, where 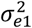 and 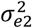 are the residual variances for site 1 and site 2, *I*_*n*1_ and *I*_*n*2_ are identity matrices, n1 and n2 are the number of individuals in each site. When we estimated the heritability for joint-sites, variance structures for family by site interaction in *μ* and *e* were assumed as homogeneous and family effect was added in the model in order to dissect G×E.

### Statistical analyses for genomic predictions

GBLUP, Bayesian ridge regression (BRR), BLASSO, and reproducing kernel Hilbert space (RKHS) were used to estimate GEBV. We implemented GBLUP calculations using ASReml R [36]. And we implemented the BRR, BLASSO, and RKHS methods using BGLR function from the BGLR package in R [37]. The details of these statistical methods will be defined later. The GEBVs were estimated using the following mixed linear model:

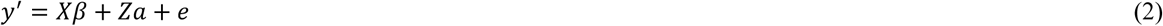

Where *y*’ is a vector of adjusted phenotypic observation by post-block effect and standardized site effect (transforming it to have zero mean and unit variance for each site), *β* is a vector of fixed effect, including a grand mean), *a* and *e* are vectors of random additive and random error effects, respectively, and *X* and *Z* are the incidence matrices.

The traditional pedigree-based best linear unbiased prediction (ABLUP) was used to compare with the four genomic-based best linear unbiased prediction methods.

#### 1) ABLUP

The ABLUP is the traditional method that utilizes a pedigree relationship matrix (A) to predict the EBV. For ABLUP the vector of random additive effect (a) in equation (2) is assumed to follow a normal distribution 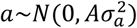, where 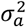 is the additive genetic variance. The residual vector e is assumed as 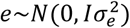, where *I* is the identity matrix. The mixed model equation (2) was solved to obtain EBV as:

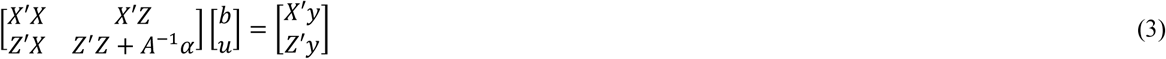

The scalar α is defined as 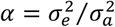, where 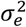 is the residual variance, 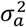 is the additive genetic variance.

#### 2) GBLUP

The GBLUP model is the same as ABLUP, with the only difference being that the genomic relationship matrix (G) replaces the A matrix. The G matrix is calculated as 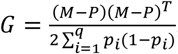, where *M* is the matrix of samples with SNPs encoded as 0, 1, 2 ( i.e. the number of minor alleles), *P* is the matrix of allele frequencies with the *i*^th^ column given by 2(*p*_*i*_ − 0.5), where *p*_*i*_ is the observed allele frequency of all genotyped samples. In GBLUP, the random additive effect (a) in equation (2) is assumed to follow 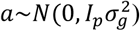, where 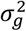 is the genomic-based genetic variance and GEBVs 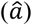 are predicted from equation (3), but with A^-1^ replaced by G^-1^ and 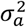 replaced by 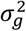. The inverse of G matrix was estimated using write.realtionshipMatrix function in synbreed package in R.

#### 3) Bayesian ridge regression (BRR)

BRR is a Bayesian version of ridge regression with the shrinkage merit that was originally intended to deal with the problem of high correlation among predictors in linear regression models [38]. The random additive vector *a* is assigned a multivariate normal prior distribution with a common variance to all marker effects, that is 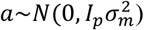, where *p* is the number of markers. Parameter 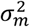 denotes the unknown genetic variance contributed by each individual marker and is assigned as 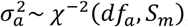, where *df*_*a*_ are degrees of freedom, *S*_*m*_ is the scale parameter. Finally, the residual variance is assigned as 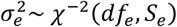, where *df*_*e*_ is degrees of freedom for residual variance, *S*_*e*_ is the scale parameter for residual variance.

#### 4) Bayesian LASSO (BLASSO)

BLASSO is a Bayesian version of LASSO regression with two properties of LASSO: 1) shrinkage and 2) variable selection. BLASSO assumes that the random additive effects in equation (2) is given a multivariate normal distribution with marker specific prior variance, which is assigned as 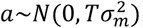, where 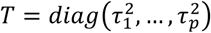, Parameter 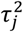 is assigned as 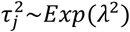 for j=1,…, *p*, where *λ*^2^ is assigned as *λ*^2^~*Gamma*(*r, δ*). The residual variance is assigned as 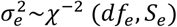, where *df*_*e*_ is degrees of freedom, *S*_*e*_ is the a scale parameter.

#### 5) RKHS

RKHS assumes that the random additive marker effects in equation (2) are distributed as 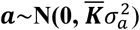, where 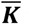 is computed by means of a Gaussian Kernel that is given by *K*_*ij*_ = exp(−*hd*_*ij*_), where *h* is a semi-parameter that controls how fast the prior covariance function declines as genetic distance increase and *d*_*ij*_ is the genetic distance between two samples computed as 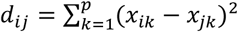, where *x*_*ik*_ and *x*_*jk*_ are the *k*^th^ SNPs for the *i*^th^ and *j*^th^ samples, respectively [39]. RKHS method uses a Gibbs sampler for the Bayesian framework and assigned the prior distribution of 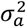 and 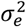 as 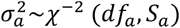 and 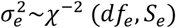, respectively. Here we chose a Single-Kernel model as suggested by Perez and de Los Campos [37], where h value was defined as h=0.25.

### Model convergence and prior sensitivity analysis

The algorithm is extended by Gibbs sampling for estimation of variance components. The Gibbs sampler was run for 150,000 iterations with a burn-in of 50,000 iterations. A thinning interval was set to 1000. The convergence of the posterior distribution was verified using trace plots. Flat priors were given to all the models.

### Heritability and type-B genetic correlation estimates

Pedigree-based individual narrow-sense heritabilities 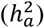 and marker-based individual narrow-sense heritabilities 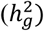 were calculated as

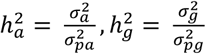

respectively, where 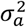 is the pedigree-based additive variance estimated from ABLUP, while 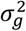 is the marker-based additive variance estimated from GBLUP. 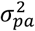 and 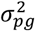 are phenotypic variances for pedigree-based and marker-based models, respectively. Type-B genetic correlation was calculated as 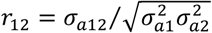, where *σ*_*a*12_ is covariance between additive effect of the same traits in different sites and 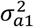 and 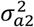 are estimated additive variance for the same traits in different sites, respectively [40].

### Model validation and estimation of GS accuracy

Tenfold cross-validation (90% of training and 10% validation) was performed for all the models, except in testing the various sizes of training data sets and the number of trees per family. GEBVs in VS for Bayesian and RKHS methods were estimated as

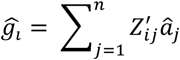

Where 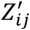 is the indicator covariate (−1, 0, or 1) for the i^th^ tree at the jth locus and 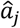 is the estimated effect at the j^th^ locus. In this study, prediction accuracy (accuracy) was defined as the Pearson correlation between cross-validated GEBV and EBV as “true” or reference breeding values estimated from ABLUP using all the phenotypic data (*y*). Predictive ability (PA) was defined as the Pearson correlation between GEBV and adjusted phenotypic value *y*’ in equation (2).

### Testing different statistical models and the size of training set on GS accuracy

To test the effect of different statistical models, we used the five models (ABLUP, GBLUP, BRR, BLASSO, and RKHS) and five TS/VS size (ratio of 1:1, 3:1, 5:1, 7:1, and 9:1 of training/validation population size, respectively) to evaluate the accuracies of different models. For each of 25 models, 10 replicate runs were carried out for each scenario of four traits.

### Testing population size (number of families) on GS accuracy

To test the effect of the number of families (population size) on GS, we randomly selected family numbers from 10 to 120 to test the efficiency of GS. The purpose is to examine efficiency of using a small subset of families in clonal selection.

### Testing family size (number of trees per family) on GS accuracy

To test the effect of the number of trees per family on GS, we randomly selected 1 to 20 trees per family as TS, remaining trees as VS.

### Testing site effect on GS accuracy

In order to consider genotype and environment interaction (G×E) effect on GS, different GS scenarios were tested: 1) within-site GS, both training and validation sets are from one single site, where EBV from single site model including G×E term was assumed as true breeding value of the site; 2) cross-site GS-using one site data as TS to predict GEBV in another site. 3) joint-site GS-average EBV was assumed as true breeding value.

### Testing the effect of relatedness on GS accuracy

In order to test whether the different relatedness (family structures) affect the accuracy and PA, three different scenarios were used in this study. There were: 1) TS and VS selected based on random sampling, but more likely from the same full-sib families; 2) for comparison purpose, TS and VS selected based on half-sib family structure in which the TS and VS shared female parents, but from different families; 3) for comparison purpose, TS and VS selected based on unrelated family structure in which the TS and VS shared different female and male parents.

### Testing subset of markers (SNPs) on GS accuracy

To test the impact of the number of SNPs on the accuracy of genomic prediction models, 15 subsets of SNPs (10, 25, 50, 100, 250, 500, 750, 1K, 2K, 4K, 6K, 8K, 10K, 50K, and 100K) and two different types of sampling strategies: 1) randomly selected SNP subsets and 2) SNP subsets selected with the largest positive effects were implemented. These were done using full-sib and half-sib families to examine the effect of relatedness on the number of markers required for effective GS selection. Single-marker regression for association testing was conducted to obtain single maker effects.

### Selection response with genomic selection

Selection response could be calculated as the ratio between selection accuracy and breeding cycle length in years.

The relative efficiency of GS to TS is

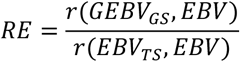

Thus, the relative efficiency of GS to TS per year is

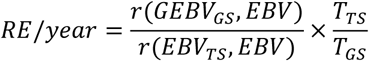

Where EBV is the estimated breeding value using the full data from model [1], and *T*_*TS*_ and *T*_*GS*_ are the breeding cycle lengths under TS and GS, respectively [16]. We assumed that the T_GS_ is reduced to 12.5 years from the current approximately 25 years of a breeding cycle by omitting or reducing progeny testing time about 10-15 years.

## Results

### Heritability and type-B genetic correlation

Heritabilities of the tree height based on GBLUP from two single sites and the joint sites were higher than those from ABLUP models (Table 1). This is in contrast with the wood quality traits (Pilodyn, velocity, and MOE) where all heritabilities from the GBLUP models were lower than those obtained from the ABLUP models. For example, heritability of Pilodyn (0.34) from the GBLUP model was smaller than that from the ABLUP model (0.41).

**Table 1.**
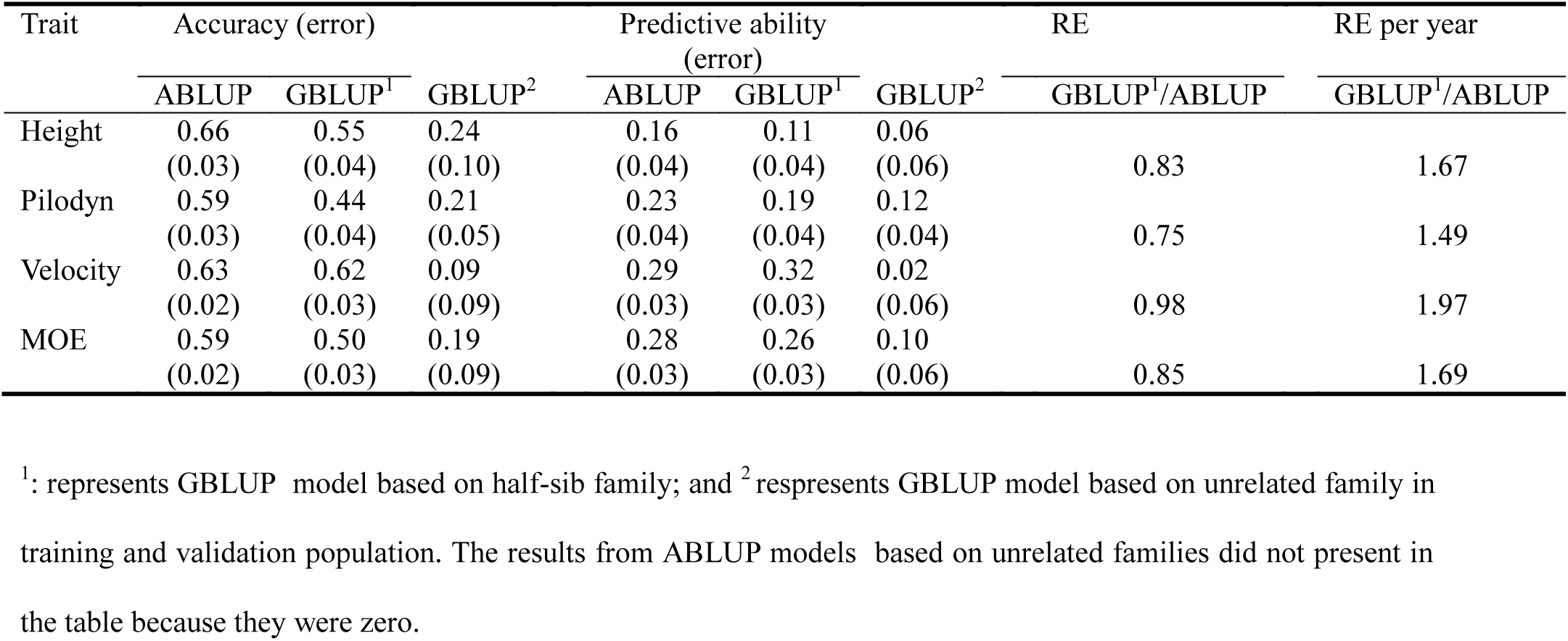
Estimates of variance components and heritabilities from the conventional pedigree-based relationship matrix model (ABLUP) and genomic-based relationship matrix model (GBLUP) in two single sites and a joint-site analysis

The type-B genetic correlations for tree height estimated from ABLUP and GBLUP models (as 0.48 and 0.41, respectively), were significantly different from 1, however, the type-B genetic correlations of wood quality traits (0.80−0.94) were higher and statistically were not significantly different from 1.

### Accuracy of different statistical methods and the size of training set

Estimates of accuracy were obtained using different statistical methods and for different ratios of TS/VS for each of four traits (Fig. 1). It was observed that ABLUP had higher accuracy than that using the four genomic selection methods (GBLUP, BRR, BLASSO, and RKHS) for tree height, Pilodyn, velocity, and MOE. Tree height had higher accuracy than the three wood quality traits using ABLUP and GS (Fig. 1 and Table 2). Among the four GS methods, GBLUP, BLASSO, and RKHS had approximately similar accuracies, but a slightly higher accuracy was observed when BRR was implemented for tree height. Nevertheless, these four GS methods had little differences on accuracy for the three wood quality traits.

**Table 2.** Accuracy, predictive ability (PA), relative efficiency (RE), and relative efficiency per year (RE per year) based on all the markers and five genomic selection scenarios for height, Pilodyn, velocity, and MOE.

| Trait | GS scenario | Accuracy (Error) |  | Predictive ability (Error) |  | RE | RE per year |
| --- | --- | --- | --- | --- | --- | --- | --- |
|  |  | ABLUP | GBLUP | ABLUP | GBLUP | GBLUP/<br>ABLUP | GBLUP/<br>ABLUP |
| Height | Site1 | 0.82 (0.01) | 0.74 (0.02) | 0.15 (0.03) | 0.16 (0.04) | 0.91 | 1.81 |
|  | Site2 | 0.89 (0.01) | 0.77 (0.02) | 0.26 (0.03) | 0.23 (0.03) | 0.87 | 1.73 |
|  | Site1→2 | 0.48 (0.04) | 0.39 (0.04) | 0.10 (0.04) | 0.11 (0.03) | 0.81 | 1.63 |
|  | Site2→1 | 0.54 (0.02) | 0.48 (0.02) | 0.02 (0.03) | 0.01 (0.03) | 0.89 | 1.77 |
|  | Joint-site | 0.91(0.00) | 0.81 (0.02) | 0.20 (0.02) | 0.20 (0.02) | 0.89 | 1.78 |
| Pilodyn | Site1 | 0.69 (0.02) | 0.60 (0.03) | 0.29 (0.04) | 0.24 (0.05) | 0.87 | 1.74 |
|  | Site2 | 0.72 (0.02) | 0.58 (0.03) | 0.33 (0.03) | 0.29 (0.04) | 0.80 | 1.60 |
|  | Site1→2 | 0.57 (0.03) | 0.52 (0.03) | 0.28 (0.03) | 0.28 (0.03) | 0.92 | 1.84 |
|  | Site2→1 | 0.58 (0.02) | 0.52 (0.02) | 0.24 (0.04) | 0.23 (0.03) | 0.90 | 1.80 |
|  | Joint-site | 0.77 (0.01) | 0.66 (0.01) | 0.32 (0.01) | 0.30 (0.02) | 0.86 | 1.71 |
| Velocity | Site1 | 0.75 (0.01) | 0.70 (0.02) | 0.42 (0.03) | 0.44 (0.04) | 0.93 | 1.87 |
|  | Site2 | 0.80 (0.01) | 0.69 (0.02) | 0.41 (0.02) | 0.37 (0.03) | 0.86 | 1.72 |
|  | Site1→2 | 0.72 (0.02) | 0.63 (0.02) | 0.40 (0.03) | 0.36 (0.03) | 0.88 | 1.76 |
|  | Site2→1 | 0.69 (0.02) | 0.60(0.03) | 0.37 (0.02) | 0.34 (0.03) | 0.87 | 1.75 |
|  | Joint-site | 0.83 (0.01) | 0.74 (0.01) | 0.43 (0.02) | 0.41 (0.02) | 0.89 | 1.78 |
| MOE | Site1 | 0.70 (0.01) | 0.64 (0.03) | 0.34 (0.03) | 0.35 (0.05) | 0.91 | 1.83 |
|  | Site2 | 0.76 (0.02) | 0.64 (0.03) | 0.38 (0.04) | 0.35 (0.05) | 0.84 | 1.69 |
|  | Site1→2 | 0.66 (0.02) | 0.59 (0.02) | 0.34 (0.02) | 0.32 (0.03) | 0.89 | 1.78 |
|  | Site2→1 | 0.61 (0.03) | 0.58 (0.02) | 0.29 (0.04) | 0.31 (0.03) | 0.95 | 1.89 |
|  | Joint-site | 0.79 (0.01) | 0.69 (0.03) | 0.38 (0.02) | 0.36 (0.03) | 0.87 | 1.75 |

**Fig. 1.**
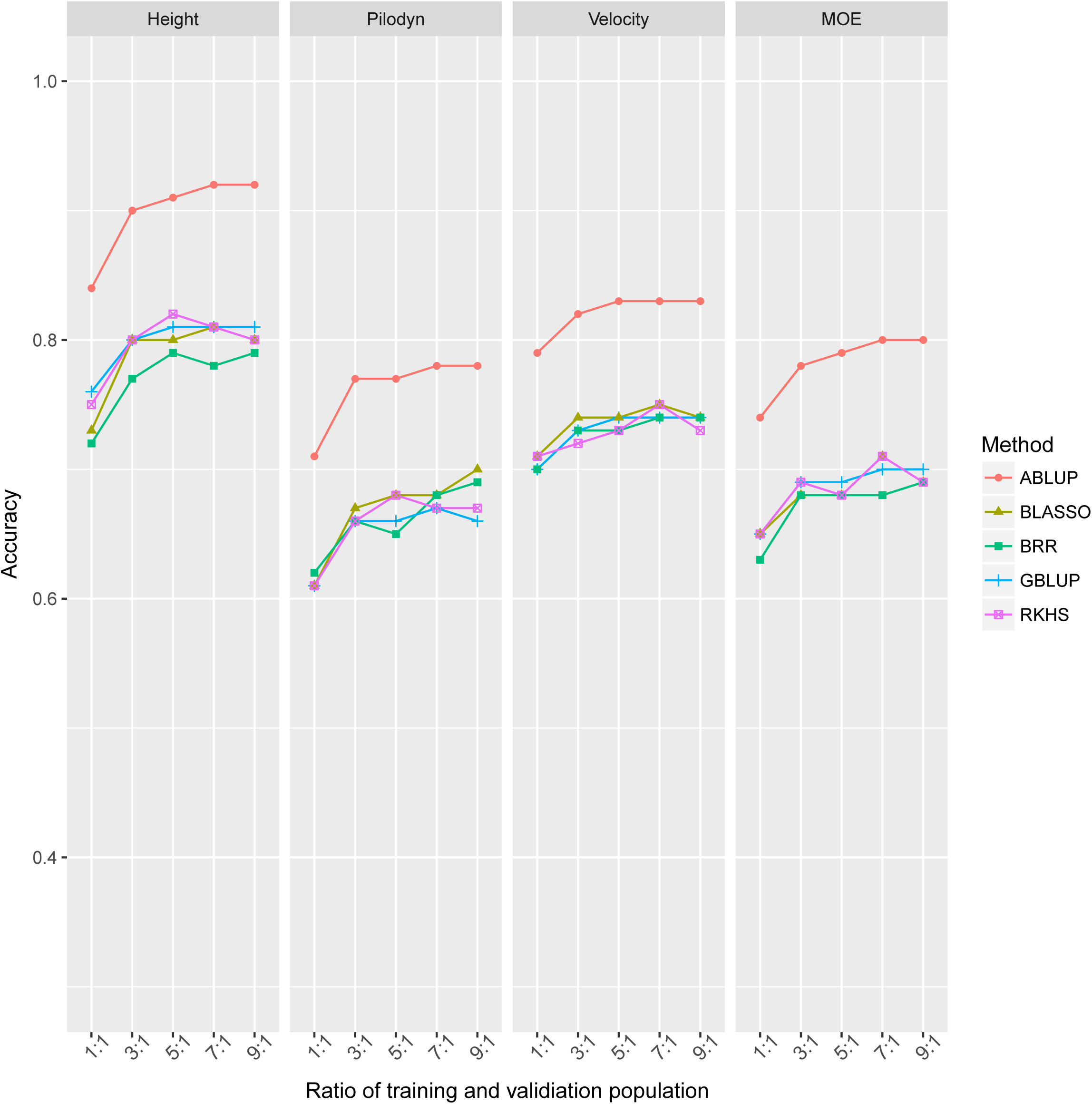
Accuracy of different methods and increasing ratios of training set (TS) and validation set (VS).

GS accuracy increased as the ratio of training to validation population size increased. However, for GS methods, the maximum accuracy was reached at the ratio of 5:1 between training to validation populations for tree height while maximum accuracy seemed to change minimally after the 3:1 ratio for wood quality traits.

### Impact of population size (number of families) on GS accuracy

We estimated the effect of population size (number of families) on GS. The accuracies of all four traits increased when the number of families increased from 10 to 120 families in both the ABLUP and GBLUP model building (Fig. 2). PA had a similar trend, except for tree height. PA of tree height increased from 10 to 30 families and then had similar values up to 120 families in model building.

**Fig. 2.**
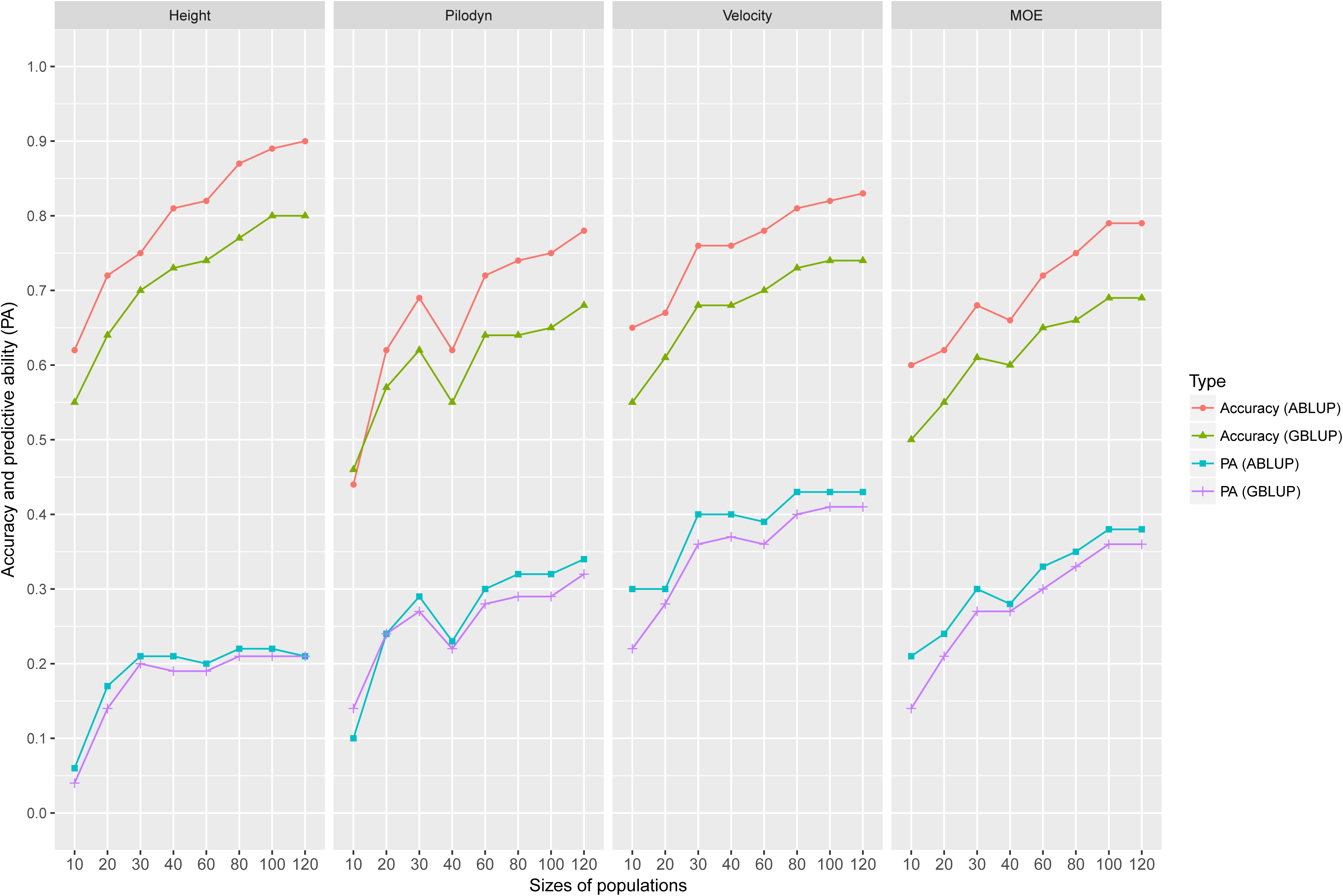
Accuracy and predictive ability (PA) of genomic selection with different sizes of population (subset of families) based on two statistical methods: 1) ABLUP and GBLUP with 9:1 for training set and validation set.

### Impact of number of trees per family on GS accuracy

In this study, all 128 full-sib families planted in both progeny trials were selected. Five trees per family in site 1 (Vindeln) and 20 trees per family were selected if there were enough trees for some families in site 2 (Hädanberg). Thus, in joint-site cross-validation model, maximum 20 trees per family were tested. Accuracies and PA from ABLUP for all the traits were higher than those from GBLUP when we randomly selected a subset trees per family as TS (Fig. 3). It was also observed that accuracy and PA had similar increased trends as the number of trees within-family increased, but it flattened (stabilized) as the tree numbers reached between 6 and 14, depending on method (e.g. ABLUP and GBLUP) and traits. The accuracies of tree height, Pilodyn, velocity, and MOE increased initially from 0.48, 0.39, 0.52, and 0.41 to 0.84, 0.64, 0.78, and 0.72, respectively, and then stabilized after tree number reached 14, 6, 12, and 12 per family for ABLUP, respectively. The GS accuracies of tree height, Pilodyn, velocity, and MOE also stabilized after the tree number reached 18, 6, 10 and 10 per families for GBLUP, respectively. This may indicate that more trees within a family (16–19) are required for a reliable training set in GS for growth trait than for wood quality traits (6–12).

**Fig. 3.**
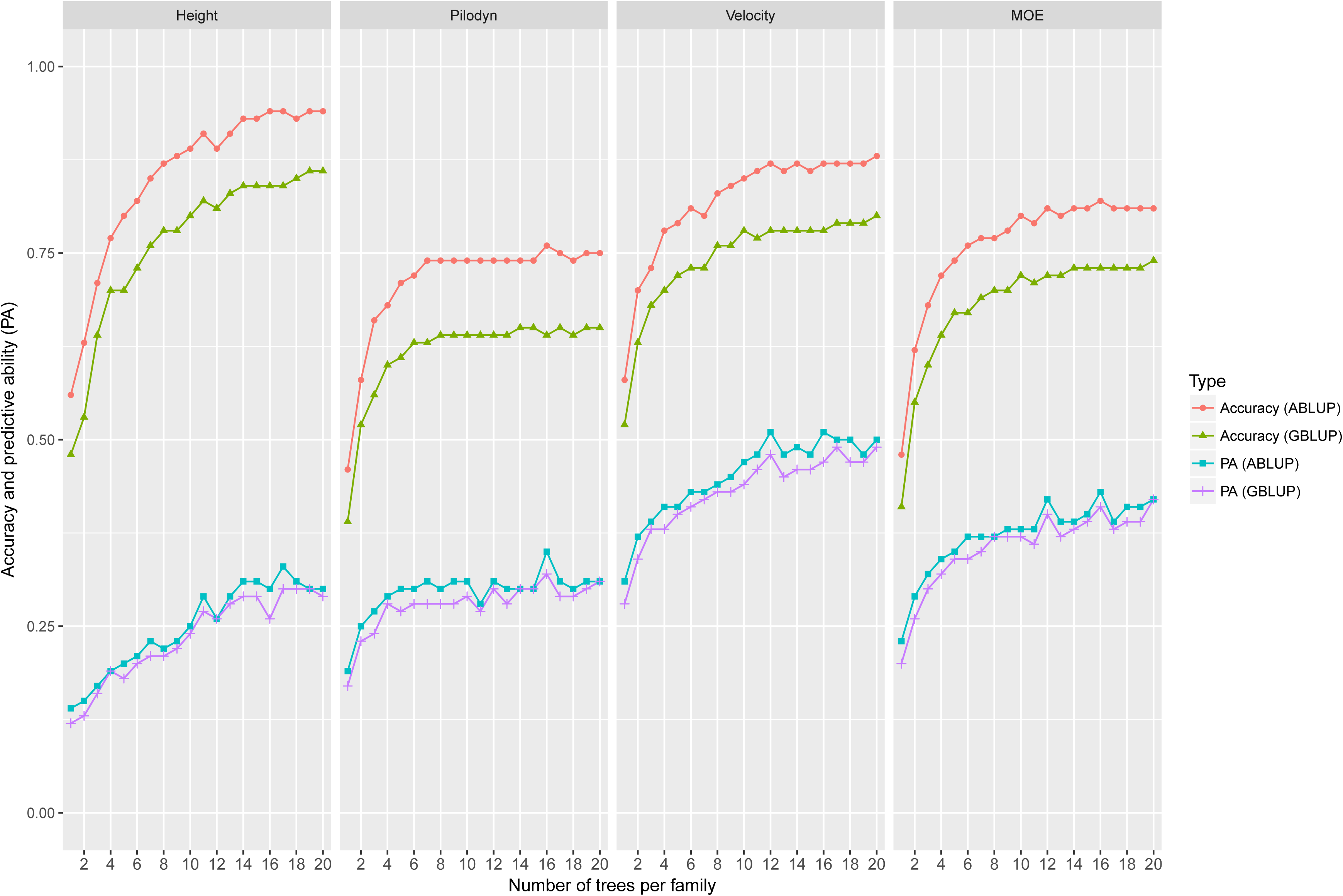
Accuracy and predictive ability (PA) of genomic selection with different subsets of trees per family based on two statistical methods: 1) ABLUP with randomly selecting subset from one to 20 trees per family as training set (TS); 2) GBLUP with randomly selecting subset from one to 20 trees per family as TS.

### Testing site effect on GS accuracy

Accuracy and PA of ABLUP and GBLUP were listed in Table 2 for three selection scenarios: Within-site training and selection, cross-site training and selection (e.g. training based on one site while selection for the second site) and joint-site training and selection using full-sib family structure. For all four traits, accuracy of within-site training and selection were always higher than the second scenario of cross-site training and selection. However, the accuracy differences between within-site and cross-site were larger for tree height than for three wood quality traits. For example, average accuracy of two within-site selections is 0.76 relative to 0.44 for average across-site for tree height from GBLUP. However, for MOE, average accuracy of two within-site selection is 0.64 relative to 0.59 for average cross-site selection. PA had a similar pattern. The joint-site model accuracies from both ABLUP and GBLUP were the highest than within-site and cross-site training and selection. Especially, for instance, tree height accuracy in the joint-site (0.81) was higher than average of within-site (0.76) and average of cross-site (0.44). Similarly, PAs from the joint-site model for all four traits were higher than cross-site training and selection model, but were not always higher than within-site training and selection model. For instance, PAs from ABLUP (0.26) and GBLUP (0.23) models in site 2 were higher than those from the joint-site (0.20 in both models) for tree height.

Relative efficiency (RE) of GBLUP to ABLUP was lower than 1 for all the selection scenarios and traits, ranging from 0.80 to 0.95 with an average of 0.88 However, RE per year (assuming halving a breeding cycle time) reached from 1.60 to 1.89 with an average of 1.76 and were not related to any traits and selection scenarios.

### Relatedness

Compared with full-sib family structure, GS models built with a half-sib family structure led to a considerable decrease in accuracy and PA (Table 3). For instance, in the half-sib family structure, GS accuracy and PA from GBLUP model decreased from 0.81 and 0.20 to 0.55 and 0.11, respectively for tree height and from 0.69 and 0.36 to 0.50 and 0.26, respectively for MOE. However, both RE and RE per year changed little between half-sib and full-sib structure in GS selection. For example, RE and RE per year increased slightly from 0.89 and 1.78 to 0.98 and 1.97, respectively for velocity from full-sib to half-sib population while RE and REs per year decreased slightly from 0.89 and 1.78 to 0.83 and 1.67, respectively from full-sib to half-sib population for tree height.

**Table 3.**
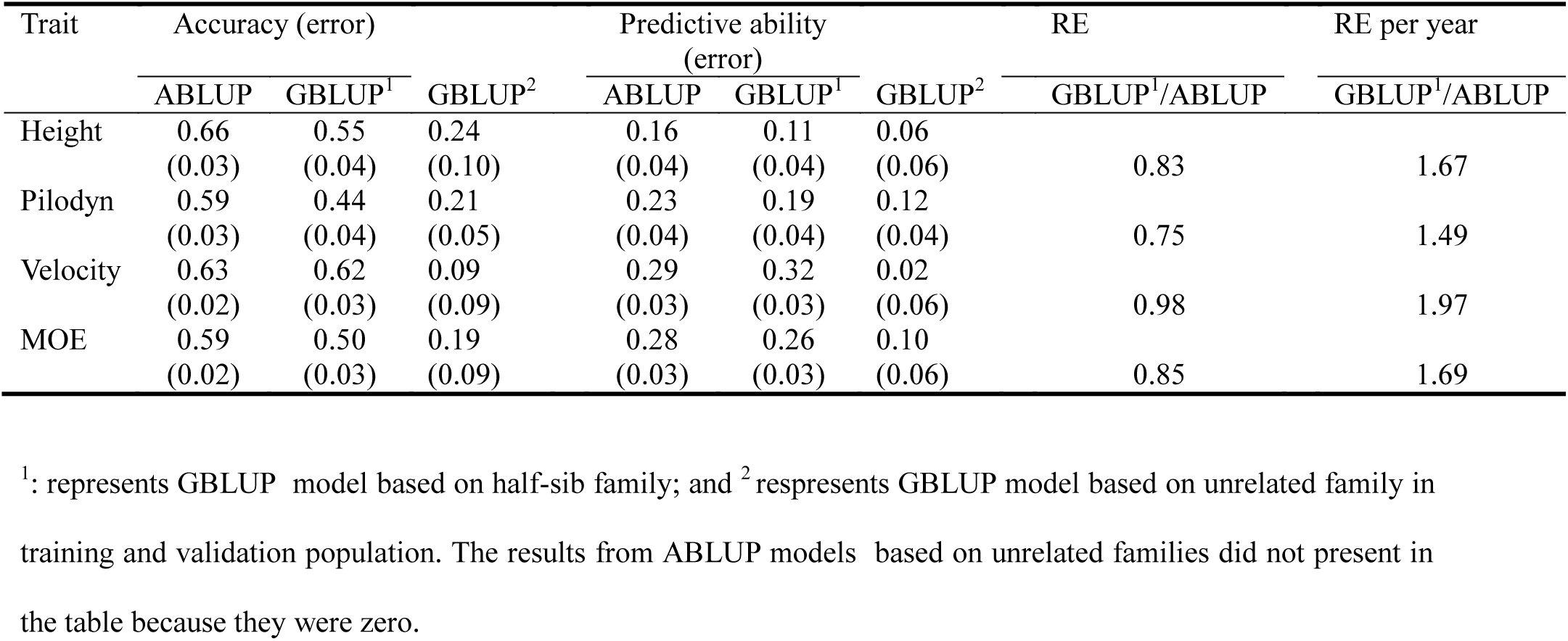
Accuracy, predictive ability (PA), relative efficiency (RE), and relative efficiency per year (RE per year) of genomic selection model based on half-sib families and unrelated families using all markers in a joint-site analysis

Compared with half-sib family structure, GS models built with unrelated family between TS and VS had a considerable decrease in accuracy and PA from GBLUP (Table 3). For example, in the unrelated family structure, GS accuracy and PA from GBLUP model decreased from 0.55 and 0.11 to 0.24 and 0.06, respectively for tree height and from 0.50 and 0.26 to 0.19 and 0.10, respectively for MOE. Especially for velocity, GS accuracy and PA from GBLUP model with a marked decrease from 0.62 and 0.32 to 0.09 and 0.02, respectively, was observed. However, it is worth to note that ABLUP models built with unrelated family had zero accuracy between training and validation populations.

### Impact of the number of SNPs on GS accuracy

Accuracy and PA using subsets of markers with the largest positive effects were higher than those using subsets of random markers until the subset of random markers reached 100K SNPs (Fig. 4). Accuracy and PA using a subset of random markers increased continuously with the increase of the number of markers, until all the markers were included. However, accuracy and PA using the subset of markers with the largest positive effects showed different trends. It increased initially, then stabilized and finally decreased until it got to the same level as the random markers selection at the highest number of markers of 100K SNPs. The use of subsets of markers with the largest positive effects had higher accuracy and PA than random markers until all the markers were used. For example, PA using a subset of markers with the largest positive effects increased initially from 0.20 to 0.40 with 250 SNPs and then decreased to 0.22 with 10K SNPs for tree height.

**Fig. 4.**
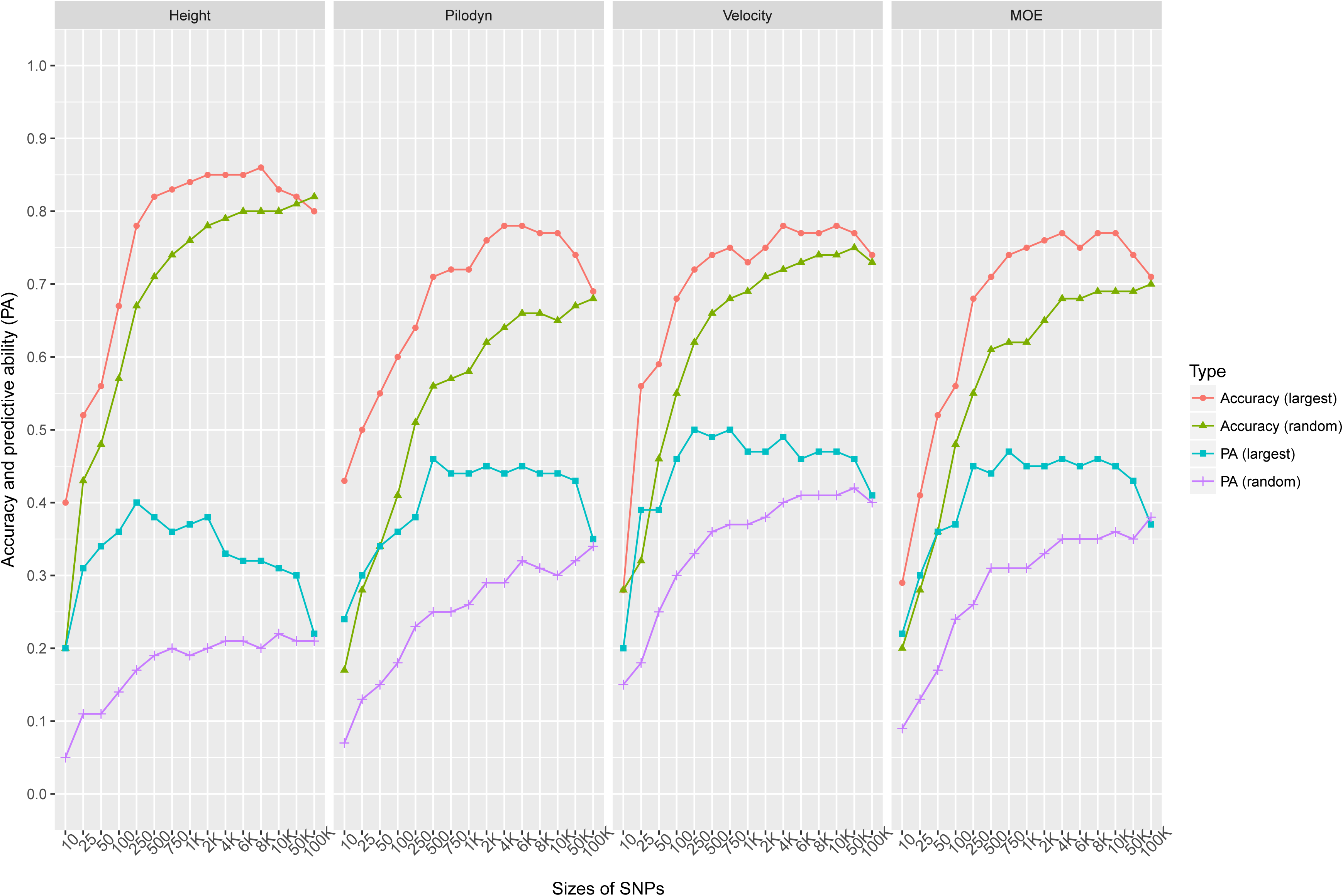
Accuracy and predictive ability (PA) of genomic selection with subset SNPs based on 2 scenarios: 1) randomly selecting the SNPs subset (10, 25, 50, 100, 250, 500, 750, 1000, 2000, 4000, 6000, 8000, 10000, and 100000 SNPs); 2) selecting the SNPs subset with the largest positive effects.

Trait also had an influence on the number of markers to reach the plateau of the accuracy and PA using a subset of markers with the highest positive effects. The accuracy and PA reached a plateau when the number of markers increased to approximately 8K for tree height while the number of markers only needed to increase to 4 to 6K for the accuracy and PA to reach a plateau for the three wood quality traits.

### Impact of the number of SNPs and relatedness on GS accuracy

Accuracy and PA using subsets of random markers with full-sib family structure were higher than that using subset of random makers with half-sib family structure (Fig. 5). To reach accuracy of 0.6 for tree height, full-sib families only needed 100 markers while half-sib families needed approximately 8K markers. Similarly, for velocity, full-sib family only needed approximately 250 markers while half-sib families needed approximately 8K markers. For the four traits, the accuracies observed with 250 markers in full-sib family structure were reached the similar values with the application of all 100K markers used in the half-sib family structure. For example, the accuracy (0.67) for tree height obtained with 250 markers was larger than that observed with any subsets of markers from 250 to 100K (0.61) for half-sib family structure. Similarly, PA with random 750 markers used in full-sib family structure reached the similar level as in half-sib family structure using all 100K markers.

**Fig. 5.**
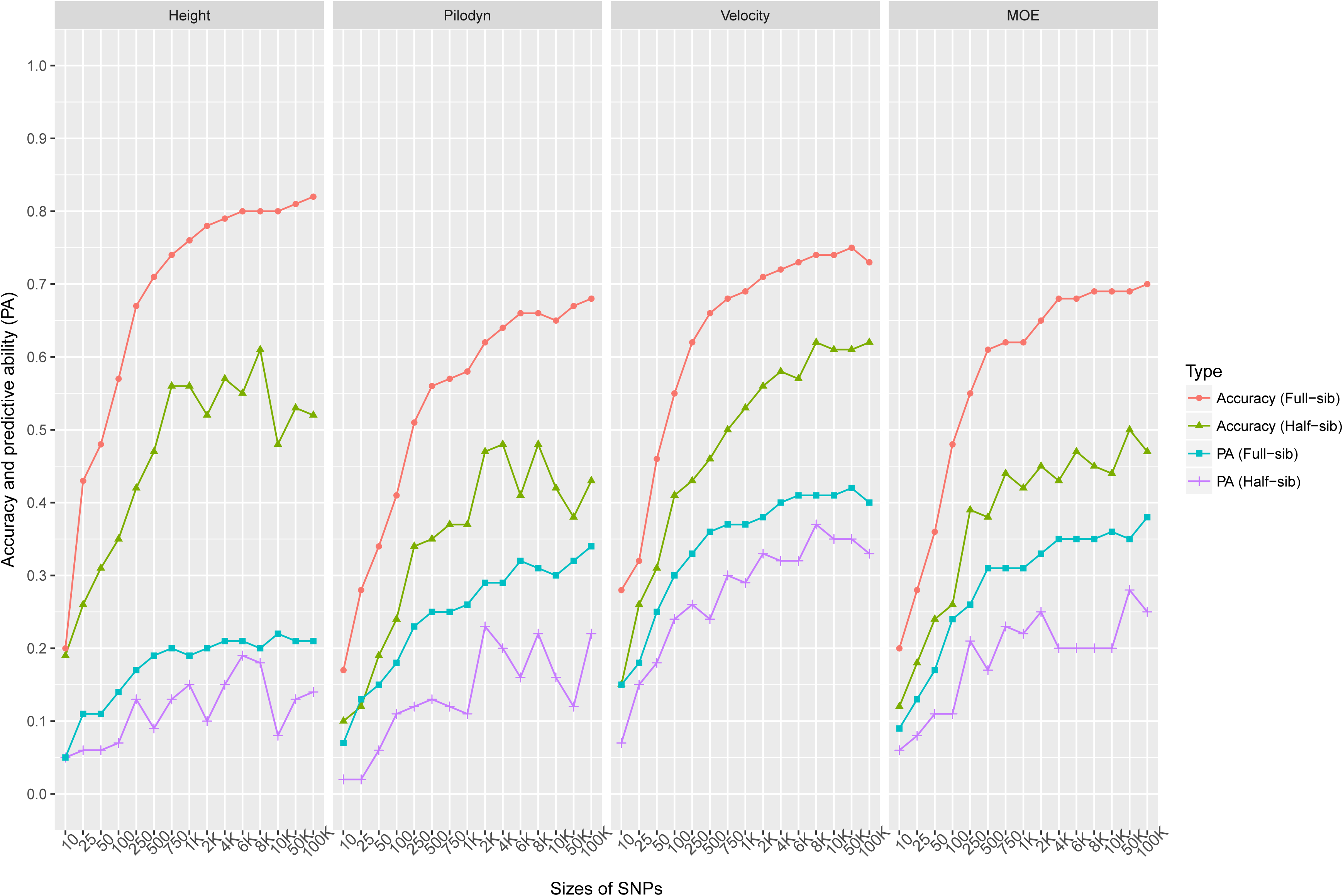
Accuracy and predictive ability (PA) of genomic selection with subset SNPs based on 2 scenarios: 1) randomly selecting subset of SNPs (10, 25, 50, 100, 250, 500, 750, 1000, 2000, 4000, 6000, 8000, 10000, and 100000 SNPs) with full-sib family structure; 2) selecting the subset of SNPs with half-sib family structure.

## Discussion

### Heritability estimates

Heritability is an essential genetic parameter in selective breeding and its values are dependent on relative contribution of genetic and environment, and varied among traits [40]. In our study, heritabilities for wood quality traits were higher than those for tree height, which is expected and agrees with previous reports for Norway spruce [1, 28]. Tan et al. [4] reported that heritability estimates obtained from GBLUP were higher than those from ABLUP for growth and wood quality traits in *Eucalyptus*. In contrast, Lenz et al. [15] and Gamal EL-Dien et al. [12] reported that heritability estimates obtained from GBLUP were lower than those from ABLUP for similar growth and wood quality traits in black spruce and interior spruce (*Picea glauca* [Moench] Voss × *Picea engelmannii* Parry ex Engelm.). In this study, the heritability estimates for tree height obtained from GBLUP were slightly higher than those from ABLUP, but there is no significant difference from ABLUP considering the standard error. The heritability estimates for wood quality traits obtained from GBLUP were slightly lower than those from ABLUP, indicating pedigree-based ABLUP model may inflate or deflate heritability estimates if estimates from GBLUP reflect true genetic relationship among families and account for Mendelian segregation within families.

The impact of heritability on the accuracy seems to be low in this study, in line with the report in Douglas-fir [19] and interior spruce [12]. In our joint-site analyses, the heritabilities of tree height, Pilodyn, velocity, and MOE were low to moderate (0.15, 0.34, 0.37, and 0.36, respectively), but the accuracies were high (0.81, 0.66, 0.74, and 0.83, respectively). Several factors may explain this: (1) the large sample size of 1370 and the relative small effective population size (*N*_*e*_=56) likely negate the effect of low trait heritability on prediction accuracy [19]. Märtens et al. (2016) proved that increasing relatedness between training and validation populations leads to high prediction accuracy on yeast; (2) the accuracy is the correlation between EBVs assuming as true breeding values from ABLUP and GEBVs from GBLUP in VS in that heritability may affect PAs of ABLUP and GBLUP, but may have little influence on the correlation; (3) the accuracy estimate only represents the additive genetic effect. We found that PA is more similar to the narrow-sense heritability because PA involved both phenotypic and genetic values. For example, heritability of MOE from GBLUP model (0.36) is the same as from ABLUP model (0.36).

In the present study, tree height accuracy (0.81 from GBLUP in Table 2) in full-sib family structure with 1233 individuals in TS was similar as in the deterministic simulation with 50 QTLs and heritability of 0.2 [16]. Wood quality traits were similar to that in the simulation with 100 QTLs and a heritability of 0.4.

### Different GS methods show similar results

As expected, the accuracies of four different genomic statistical methods did not clearly outperform each other. This contrasts with previous evaluation of the RKHS method that was reported to outperform other GS methods for low heritability growth traits (Tan et al. 2017). It was usually observed that different genomic statistical methods produce similar results for growth and wood quality traits in other forest trees species [13, 19, 41]. With an exception, Resende et al. [9] compared Ridge Regression-BLUP (RR-BLUP), Bayes A, Cπ, and BLASSO for 17 traits in loblolly pine, and found that Bayes A and Bayes Cπ have higher PA than RR-BLUP and BLASSO for fusiform rust disease-resistance trait. They attribute this to a few genes with large effects that control disease resistance. When the number of markers increases the computational time for Bayesian method took longer time to converge. Therefore, we also support the proposal that GBLUP is an effective method in providing the best compromise between computational time and prediction efficiency if there are no major gene effects [4, 42].

### The effect of training data set and number of trees per family on GS accuracy

In this study, ratio of TS/VS varied from 1:1 to 9:1. We found that the accuracy improved from the TS/VS ratios of 1:1 to 3:1, but accuracy only improved a little after the TS/VS ratio of 3:1. This is different from other studies [4, 14] that increased ratio of the TS/VS beyond 3:1 still increase the accuracy. However, we found that our result concurs with other reports when the ratio of TS/VS is related to the number of trees per family [14]. In our case, each family under 1:1 ratio of the TS/VS, has 5 trees. There is an average of 10.7 trees per each of 128 families. The ratios of TS/VS with 1:1, 3:1, 5:1, 7:1, and 9:1 equate to average number of TS trees per family of 5.3, 8.0, 8.9, 9.4, and 9.6, respectively. This may indicate that after 8 trees per family on average as TS, there is little increase of GS efficiency for full-sib family. Based on resampling technique, Perron et al. [43] reported that the number of trees per family have an important effect on the magnitude and precision of genetic parameter estimate. For tree height in that study, at least four trees per family at each site should be included in a half-sib family in order to estimate a more accurate heritability and 4−8 trees/per family per site still could improve the accuracy of heritability. However, further increase in the number of tree trees per family had little contribution toward increase of accuracy. Wood quality traits usually have higher heritabilities than those of growth traits and also have less G×E. Therefore, wood quality traits may need a lower number of trees than growth traits for obtaining a similar accurate estimate for genetic parameters. Such calculations could guide us to make a more accurate estimate on number of trees per family required for phenotyping and genotyping.

### The effect of population size (number of families) on GS accuracy

The effect of the number of families used in cross validation test was found important in this study with full-sib family structure. We found that the PA and accuracy increased greatly for all the traits from 10 to 120 families, except the PA for tree height, which stabilized after 30 families for cross validation test. Based on resampling technique, Perron et al. [43] reported that the number of families has a less important effect on the magnitude and precision of genetic parameter estimate in a half-sib family structure. While in this study, we found that the number of families is also important for estimates of GS accuracy and PA. It may be due to small effective population size (55 unrelated parents) compare to the study in Perron et al. [43]. One of application of GS is in clonal forestry to select the best clones after selection and mating of several best parents (5−10). One question is how to build the training equation for such clonal selection. Should we use progenies of the selected parents or all parents of the trial? From this study, it seems that it is more efficiency involving progenies from a larger number of parents (families). Model building and selection using 10 families seems to have lower accuracy than using a larger number of families. We don’t know whether it is due to small size of families (10−20 trees per families) used in this study, and whether increasing the family size (for example 40 trees per family) as used in clonal progeny testing in Norway spruce in Swedish tree breeding program [2] would increase GS accuracy in a small group of elite families.

### Genotype-by-environment interaction

G×E is important when the seedlings are planted in different environments. We found that tree height in these two northern trials had a significantly strong G×E, indicated by the type-B genetic correlations (0.48 and 0.41) from ABLUP and GBLUP, respectively (Table 1). Such strong G×E resulted in a low accuracy and PA when one site is used as TS to predict BVs in another site as VS. A moderate to strong G×E for growth traits has been reported in several studies in southern and central Sweden [22, 44, 45], but not documented in northern Sweden. Chen et al. [23] reported that within a test serial in southern and central Sweden, the averages of type-B genetic correlations varied from 0.60 to 0.89 in 6 test series and their type-B correlations were higher than those in this study. Such strong G×E in the present study should be considered in model fitting in order to improve PA and accuracy. Several advanced models have been built and tested in crops [46, 47] and one study in tree species [48]. For instance, Oakey et al. [46] used maker and maker by environment interaction by RR-BLUP method to extend genomic selection to multiple environments.

As expected, we found that wood quality traits have no significate type-B genetic correlations and negligible change of accuracy and PA. A similar result has been reported in two southern Norway spruce open-pollinated progeny trials [28]. All these indicate that we could use a genomic model in one site to predict GEBV in another site in the same test series for wood quality traits.

### Effect of different family structure

We found that our genomic accuracy for tree height and Pilodyn using half-sib family was lower than that reported by Lenz et al. [14] in black spruce, even though more SNPs were used in this study (116K in 20,695 contigs vs 0.49K from SNP chip). The difference is likely due to lower heritabilities (i.e. 0.15 VS 0.42 for tree height in GBLUP) and the larger *N*_*e*_ in this study (55 VS 27 unrelated parents). Accuracy and PA decreased from full-sib to half-sib family structure and from half-sib to unrelated family structure, indicating that GS model is more efficient in strong structured populations where relatedness and LD are higher, and full-sib families also needed less number of markers than half-sib families for obtaining similar accuracy (Fig. 5). Similar results were obtained by other studies [11, 14, 41, 49]. However, the relative efficiency between ABLUP and GBLUP (the accuracy ratio) is more or less similar in both full-sib and half-sib populations, which indicates that GS could be used in both half-sib and full-sib populations. The lower estimates of accuracy and PA (Table 3) obtained from GBLUP for unrelated family structure may be due to lower LD between marker and QTLs in the unrelated population.

### The effect of number of SNPs and LD in genomic prediction

To our knowledge, this study has used the largest number of SNPs (116,765 SNPs from 20,695 contigs) for GS in tree species [3, 7, 19]. When we used GS in the half-sib family structure, the accuracy and PA reduced about 20%. This may indicate that such a large number of SNPs may still not be enough to capture most of the QTL effects due to low LD. In humans, the exome constitutes a mere 1% of the whole genome (3Gb) [50]. For Norway spruce, however, the genome size is ca. 20 Gb and LD is lower than Humans [21]. Norway spruce has a mapped genome size of 3326.3 centiMorgan (cM) (Bernhardsson et al. 2018, unpublished), which is larger compared to ~2100 cM in white spruce (*Picea Glauca* (Moench) Voss) [51]. There were about 5.6 SNPs per contigs/genes on average based on 20,695 contigs/genes that our SNPs come from. This translates an average genome coverage of ~6.2 markers/cM.

Accuracy and PA obtained from the subset of markers with the largest positive effects were slightly higher than those from a subset of markers based on random selection, implying that using the subset of markers with the largest effects in genomic regions with small LD decay could track relationship more effectively than random markers. It also implies that using the subset of markers with the largest positive effect could also obtain some effects based on part of the short-range LD [14, 41]. Thus, this factor could also be potentially useful to reduce genotyping the number of SNPs and GS cost, in highly structured population when the genome location of markers is known.

### Efficiency of genomic selection

In the present study, we observed that the efficiency of GS per year is greater than that based on traditional progeny test selection if GS is used to half generation time. In traditional pedigree-based progeny selection, the generation time for Norway spruce in Northern Sweden is at least 25 years, which is based on clonal replicated progeny trial of selected breeding trees. The clonal based progeny test procedure includes: seed sowing and growing to a sufficient size (2 yrs), vegetative propagation of seed plants by rooted cuttings (2 yrs), testing in field trials (15 yrs), and assessment of trials (1 yr). The final stage, the completion a crossing scheme to create the next generation, is about 5 yrs. If we could omit the progeny testing of the first three stages (a total of 19 yrs) and complete flowering induction and mating within 15 years of GS selection, the time for a breeding cycle could be halved for Norway spruce. In this study, we only considered the efficiency of GS based on the timing of breeding cycle. If the cost for field testing is considered, the benefit of GS might be higher if the cost for establishing and maintaining 3−4 progeny trials in each breeding population is more than genotyping.

## Conclusions

As expected, the main advantage of genomic selection is the potential to shorten the breeding cycle. We observed that:

1. Using G matrix slightly altered heritability relative to A matrix with a slight increase for tree height and a decrease for wood quality traits.

2. ABLUP is about 11–14% more efficiency than GBLUP for tree height and wood quality traits, while the four GS methods (GBLUP, BRR, BLASSO, RKHS) had a similar accuracy.

3. Efficiency of GS increased from 49% to 97% among four growth and wood quality traits if GS could half generation time for a breeding cycle.

4. The GS accuracy improved from the TS/VS ratios of 1:1 to 3:1, but accuracy only improved marginally after a TS/VS ratio of 3:1.

5. Number of families and family size also had effect on GS efficiency. Wood quality traits need fewer number of trees within-family than tree height for a similar GS efficiency.

6. GS accuracy decreased from full-sib to half-sib family structure and from half-sib to unrelated family structure. Number of markers need to be increased greatly for half-sib family to have a similar efficiency to full-sib family structure.

7. GS accuracy increased as number of markers increased. The accuracy reached a plateau when the number of markers increased to approximate 8K for tree height while the number of markers only needed to increase to 4 to 6K for the accuracy to reach a plateau for the three wood quality traits.

## Abbreviations

ABLUP: Pedigree-based best linear unbiased prediction
BLASSO: Bayesian LASSO regression
BRR: Bayesian ridge regression
DP: Read depth
EBV: Estimated breeding value
GATK: Genome Analysis Toolkit
GBLUP: Genomic best linear unbiased prediction
GEBV: Genomic breeding values
G×E: Genotype-by-environment interaction
GQ: Genotype quality
GS: Genomic selection
LD: Linkage disequilibrium
MAF: Minor allele frequency
MFA: Microfibril angle
MOE: Modulus of elasticity
PA: Predictive ability
Pilodyn: Pilodyn penetration
QTL: Quantitative trait locus
RE: Relative efficiency
RKHS: Reproducing Kernel Hilbert Space
SNP: Single nucleotide polymorphism
TS: Training set
Velocity: Acoustic velocity
VQSR: Variant quality score recalibration
VS: Validation set;

## Acknowledgements

The computations were performed on resources by the Swedish National Infrastructure for Computing (SNIC) at UPPMAX and HPC2N. We thank Dr. Junjie Zhang, Tianyi Liu, and Ms Xinyu Chen, Linghua Zhou for help of the DNA extraction and field assistance and Anders Fries for field work.

## Availability of data and materials

The datasets supporting the conclusions of this article are available upon request.

## Authors’ contributions

ZQC designed sampling strategy, coordinated field sampling, analyzed data, and drafted the manuscript. BAG, BK, and JW participated selection of the breeding populations, providing access to field experiments, tree height data and edited the manuscript. JB, JP, and MRGG participated collection of phenotypic data, extraction of the DNA, SNP calling, and editing of the manuscript. HXW conceived and designed the study, and assisted writing of the manuscript. All authors read and approved the final manuscript.

## Ethics approval and consent to participate

The plant materials analyzed for this study comes from common garden experiments (Plantation and clonal archives) that were established and maintained by the Forestry Research Institute of Sweden (Skogforsk) for breeding selections and research purposes. Three tree breeders in Sweden were coauthors in this paper. They agreed to access the materials.

## Consent for publication

Not applicable

## Author details

^1^Umeå Plant Science Centre, Department Forest Genetics and Plant Physiology, Swedish University of Agricultural Sciences, SE-90183 Umeå, Sweden. ^2^Skogforsk, Ekebo 2250, SE-268 90 Svalöv, Sweden. ^3^Skogforsk, Sävar 2250, SE-268 90, Sweden. ^4^CSIRO NRCA, Black Mountain Laboratory, Canberra, ACT 2601, Australia.

